# The effects of prefrontal tDCS on working memory associate with the magnitude of the individual electric field in the brain

**DOI:** 10.1101/2023.06.13.544810

**Authors:** Lais B. Razza, Stefanie De Smet, Sybren Van Hoornweder, Sara De Witte, Matthias S. Luethi, Chris Baeken, Andre R. Brunoni, Marie-Anne Vanderhasselt

## Abstract

Transcranial direct current stimulation (tDCS) over the prefrontal cortex has the potential to enhance working memory by means of a weak direct current applied to the scalp. However, its effects are highly variable and possibly dependent on individual variability in cortical architecture and head anatomy. Unveiling sources of heterogeneity might improve fundamental and clinical application of tDCS in the field. Therefore, we investigated sources of tDCS variability of prefrontal 1.5mA tDCS, 3mA tDCS and sham tDCS in 40 participants (67.5% women, mean age 24.7 years) by associating simulated electric field (E-field) magnitude in brain regions of interest (dorsolateral prefrontal cortex, anterior cingulate cortex (ACC) and subgenual ACC) and working memory performance. Emotional and non-emotional 3-back paradigms were used. In the tDCS protocol analysis, effects were only significant for the 3mA group, and only for the emotional tasks. In the individual E-field magnitude analysis, faster responses in non-emotional, but not in the emotional task, were associated with stronger E-fields in all brain regions of interest. A follow-up analysis showed that people with higher (vs. lower) E-fields magnitude in the left DLPFC were faster in the both tasks, and more accurate in the emotional task. Concluding, individual E-field distribution might explain part of the variability of prefrontal tDCS effects on working memory performance and in clinical samples. Our results suggest that tDCS effects can be more consistent or improved by applying personalizing current intensity, although this hypothesis should be confirmed by further studies.

## 1. Introduction

The prefrontal cortex and related brain areas regulate complex emotional and cognitive processes (Spielberg et al., 2008). Its abnormal activation has been associated with psychiatric disorders and cognitive impairment (Koenigs & Grafman, 2009; Zhang et al., 2022).

The activity of this brain region can be modulated via transcranial direct current stimulation (tDCS) using electrodes on the scalp that create a low-intensity electric (E-)field in the cortex which modulates neuronal excitability. This together with the favorable safety profile of tDCS applied over the prefrontal cortex has made the technique an attractive modality to ameliorate depressive symptoms and enhance cognitive performance (Majdi et al., 2022). Although promising, studies show that the prefrontal tDCS effects are mixed (Razza et al., 2020).

It has been hypothesized that the heterogeneity of prefrontal tDCS effects might be explained by the variability of E-field magnitude that penetrates into the brain. Indeed, E-field magnitude can vary across persons due to anatomical idiosyncrasies (i.e. skull thickness, cortical thickness and cortical architecture). Previous studies have aimed to investigate this question by evaluating the simulated E-fields in brain regions of interest in subsamples of depressive and older individuals, presenting a direct association between E-field and tDCS effects (Albizu et al., 2020; Indahlastari et al., 2021; Suen et al., 2020). Yet, evidence regarding this topic remains at a preliminary stage, with no studies having systematically investigated the magnitude of E-field of distinct intensities of tDCS (compared to sham) as a predictor of response in younger adults, which constitute the predominant population in tDCS trials.

Therefore, the purpose of this study was to unravel whether the effects of distinct prefrontal tDCS protocols (1.5mA, 3mA, and sham) on working memory paradigms depend on the induced individual E-field magnitude in brain regions of interest proxied by computational modeling analysis. Two working memory tasks were employed (emotional and non-emotional), both of which engage the DLPFC and recruit different brain regions, such as the anterior cingulate cortex (ACC) and subgenual ACC (sgACC). A linear and direct association between working memory performance and E-fields magnitude in the regions of interest was hypothesized. Moreover, we expected that people with higher E-field in deeper brain regions (ACC and sgACC) would present increased performance in the emotional paradigm since those regions are directly involved with emotional processes (Etkin et al., 2011).

## 2. Material and Methods

### 2.1 Design

This study was conducted at the Ghent University Hospital, Ghent University, from March 2022 to March 2023 and was approved by the Ethics Committee of the Ghent University Hospital (B6702021000839). A double-blind, sham-controlled, randomized, within-subjects trial was employed in which participants received sham tDCS, 1.5mA tDCS and 3.0mA tDCS. There was a two-week interval between sessions to ensure the elimination of carryover effects and to reduce learning effects in the cognitive tasks (Dedoncker et al., 2016)

### 2.2 Participants

The sample size was calculated using G*Power 3.1 software considering a repeated-measure ANOVA and a within-subject design. A low to moderate effect size (f=0.22), alpha of 0.05 and a power of 0.80 were applied, which generated a sample size of 35 participants. Therefore, 40 participants were included to account for potential drop-outs and data loss.

Individuals between 18 and 45 years without prior or present mental or neurological disorders were included. All individuals were screened for past or current psychiatric diagnoses (Pinninti et al., 2003) based on the Mini International Neuropsychiatric Interview (Pinninti et al., 2003; Wörsching et al., 2017) and the Beck Depression Inventory (Beck et al., 1961). Exclusion criteria were: (1) any contraindications for tDCS and/or MRI, (2) habitual smoking or abuse/dependence on other drugs; (3) pregnancy. Individuals included in this study have participated in previous studies of our lab. All participants provided written informed consent prior to participation.

### 2.3 Experimental Procedure

Participants visited the hospital on four occasions. For the first visit, an anatomical 3-Tesla MRI acquisition was acquired (T1- and T2-weighted sequences; repetition time = 1900 milliseconds, echo time = 2.2 milliseconds, flip angle = 9°, 176 slices/volume, slice thickness = 0.8 mm). In the following visits, participants underwent three experimental sessions in a controlled environment (Sup. Material - Appendix 1). In the first experimental session, a neuronavigation procedure (Brainsight, Rogue Resolutions, Inc) was performed in order to individually determine the targeted DLPFCs.

Throughout all experimental sessions, self-reported measurements were collected (Sup. Material - Appendix 1). During the tDCS, a 0-back task was performed for 10 minutes to standardize ongoing neural activity. For that, a word appeared on the computer screen and participants had to press a specific letter on the keyboard at its occurrence or a different key for any other word. After tDCS stimulation, an adverse effects scale was applied, followed by the 3- back tasks.

### 2.4 tDCS

TDCS was delivered for 20 minutes using two 25 cm² (5×5cm) rubber electrodes (DC-Stimulator Plus, NeuroConn, Germany) that were applied to the scalp using a conductive gel (Ten20®). The anode and cathode were respectively placed over the left and right neuronavigated DLPFCs to optimally target the sgACC (left: x = -38, y = +44, y = +26; right: x = 38, y = +44, y = +26) (Blumberger et al., 2018; Fox et al., 2012). These targets were marked on an individual cap and used in all further sessions. The active tDCS sessions were performed using currents of either 1.5mA and 3mA. The sham protocol was identical but consisted of a brief active period of 30 seconds fade-in and 30 seconds fade-out at the beginning and the end of the session with a current intensity of 3 mA (De Smet et al., 2021). The tDCS device allows for double-blinding as it has a ‘study mode’ that delivers both active or sham current based on a randomized imputed code. The codes were randomized via https://www.randomizer.org and were managed by a person not involved in data acquisition.

### 2.5 N-back Tasks

Non-emotional and emotional paradigms were applied using a 3-back approach (Sup. Material - Appendix 1). Throughout the three experimental sessions, the same task set-up was used, although stimuli were presented randomly. Both tasks started with instructions and consisted of three experimental blocks containing 50 stimuli each. The order of the tasks was counterbalanced. Accuracy and reaction times were assessed as outcome measures.

For the non-emotional task, alphabetic letters (A to L) were presented individually and randomly on the computer monitor for a period of 500 ms, with a 2000 ms inter stimulus interval. Participants were instructed to remember the letters. Via button presses on the keyboard (‘J’ to ‘yes’ and ‘N’ to ‘no’), they had to indicate whether the currently displayed letter on the monitor was the same as the one presented *3 trials before* or not. For the emotional task, the same parameters were employed, although a series of words obtained from a normative database (Moors et al., 2013), were presented on the screen. These words had different emotional negative and positive valences that were displayed an equal amount of times (Pe et al., 2013). Participants were instructed to remember the *emotional valence* of each word and indicate whether the valence of the current word shown was the same as presented 3-blocks before (pressing letter ‘J’) or not (pressing letter ‘N’).

A practice for both protocols was conducted priori. It contained the same parameters and rules of both n-back tasks, and finished when participants reached 60% accuracy. The order of the n-back paradigms was also counterbalanced in the practice. Via this method, participants were always working at their peak performance level, and learning effects (which typically arise in within-subjects protocols) could be attenuated.

### 2.6 Self-report measurements

The State-Trait-Anxiety Inventory (STAI) state inventory was administered at the beginning of each session (Spielberger et al., 1970) to identify the baseline psychological state of each participant. Moreover, the Visual Analogue Scale (VAS) was collected three times (at baseline, after tDCS and after the N-back tasks). Participants were asked to indicate from 0 to 100 (‘‘I do not experience this at all”, ‘‘I experience this very much”, respectively) how much they were feeling the following moods: ‘angry’, ‘tense’, ‘sad’ ‘happy’, ‘stressed’ and ‘anxious’. Higher scores reflect higher levels of perceived stress or negative affect.

### 2.7 Adverse effects scale

Adverse effects were assessed at the end of each tDCS session. Adverse effects were separated into 14 different categories: local pain, headache, neck pain, itching sensation, scratching, tingling sensation on the scalp, burning sensations under the electrodes, skins redness under the electrode, somnolence, concentration changes, fatigue, nausea, dizziness and seizure, which were based on a structured scale (Aparício et al., 2016). The adverse effects were reported in a binomial form, as participants indicated if they felt the specific adverse effect (‘yes’) or not (‘no’).

### 2.8 Computational Modeling Analysis

E-field modeling was performed using SimNIBS (Version 4.0, Copenhagen, Denmark) (Saturnino et al., 2019), a free and open-source software package that allows approximation of the tDCS-induced E-field distribution in the individual brain. First, head models were created using the *charm* routine based on individual T1- and T2-weighted structural magnetic resonance data (Puonti et al., 2020). In total, *charm* segments nine tissue types based on the provided structural MRI scan. Default conductivity values for each tissue were applied. Afterwards, the software creates a 3D tetrahedral mesh structure of each segmented tissue, which allows for the simulation of the E-field induced in each participant’s brain. We manually verified the *charm* segmentation to check for errors.

The resulting tetrahedral head mesh of each participant was used to simulate the E-field distribution resulting from the two active tDCS protocols (1.5 mA and 3 mA). Electrodes were placed over the MNI coordinates retrieved from neuronavigation, pointing towards Cz, with a thickness of 5 mm (electrode + conductivity paste). Of note, while E-field magnitudes scale linearly with intensity (e.g., the E-field magnitude of 3mA tDCS is twice the 1.5mA), its effects on the working memory performance are not necessarily linear.

Next, we extracted the E-field magnitude within the predefined brain regions of interest. These regions were the bilateral DLPFC, bilateral ACC and the bilateral sgACC (Miró-Padilla et al., 2019). The DLPFC was extracted from the Sallet atlas (Sallet et al., 2013) and the ACC and sgACC were extracted from the Brainnetome atlas (Fan et al., 2016)(Sup. Material - Appendix 2). Two different volumetric atlases were used for the sake of consistency with previous research (Bulubas et al., 2019; Suen et al., 2020).

Values analyzed in this study were the mean magnitude of the electric field (magn-E component), which represents the vector strength but not its direction. The peak E-field strength, quantified as the 99th percentile E-field magnitude, was visually inspected and is reported in the results section (Van Hoornweder et al., 2023).

## 3. Statistical Analysis

Two different statistical analyses approaches were used. One approach aimed to unveil the effects of different tDCS protocols on working memory, whereas the other aimed to unveil the relationship between E-field magnitude and working memory performance.

Statistics were performed using R software (Rstudio, version 4.2.2). For the 3-back task, missed responses and outliers were excluded from the analyses. Outliers were detected using the ‘DoubleMAD’ detector (Rosenmai, 2013), with performances lower and over three standard deviations around the median per groups being excluded, representing 5.17% and 2.7% of the total data for the non-emotional and emotional tasks, respectively.

Firstly, reaction time (in milliseconds) and accuracy (binary outcome) measures were the dependent variables, whereas *Protocol* (three levels: sham; 1.5mA; 3mA) was the independent variable. Reaction time analyses were performed using accurate responses only. Due to the positively skewed nature of reaction time data, several generalized linear mixed models (GLMM; ‘lme4’ package) were tested. The best-fitting model was selected by inspecting the distribution of residuals (via qq-plot) and checking the Akaike information criterion for comparisons. A GLMM of the gamma distribution with an identity link was retained (Sup. Material - Appendix 3). For accuracy, a binomial family was applied, with the variable ‘session’ considered as a fixed factor and ‘subject’ considered the random intercept. Pairwise analyses were performed using the ‘emmeans()’ function. For the VAS, a three (time: baseline, after-tDCS and after-N-back) by three (tDCS protocols) LMM was fitted, using ‘session’ as a fixed factor and ‘subject’ as a random intercept.

Secondly, we evaluated whether working memory performance was associated with the mean individual E-field magnitude induced by tDCS in each brain region of interest. For these analyses, the working memory performance was calculated based on a delta score of the active protocols minus performance in the sham group (i.e., Δreaction time = reaction time 3mA tDCS - reaction time of sham). The outcome measures were associated with the mean E-field via LMMs, taking ‘subject’ as a random intercept. Both the 1.5mA and 3mA tDCS were pooled into a single dataset as our main aim was to investigate inter-individual variability in the overall tDCS effects, but not differences of E-field magnitude between 1.5mA and 3mA. This separation is present in the graphical results only for means of visualization. Multiple comparison corrections were performed using the false discovery rate (‘stats’ package), which was applied separately for each outcome type and 3-back condition. Only the corrected p-values are presented Sup. Material - Appendix 4). Afterwards, a follow-up analysis was performed based on significant results of a previous study (Caulfield et al., 2022a). For this analysis participants were separated into two groups - above and below median E-field magnitude in the left DLPFC - and were associated with working memory performance by using linear regression. Finally, tolerability analysis was calculated using LMMs (Sup. Material - Appendix 4).

For all statistical tests, the significance level was set to alpha = 0.05.

## Results

A total of 40 participants were included (67.5% women, mean age 24.7 (*SD*= 5.6)). Two participants dropped out after the first session. Therefore, a total of 117 tDCS sessions (1.5mA: *n* = 39; 3mA: *n* = 38; sham: *n* = 39) was performed. Both STAI and VAS measures results did not show any significant differences among groups and timepoints (Details can be found in the Sup. Material - Appendix 5). The overall performance per tDCS protocol can be found in the Sup. Material - Appendix 6.

### tDCS protocol analysis

For the non-emotional task, no significant effect of tDCS protocols for reaction time (chi-squared (χ2)(2) = 3.12, p = 0.21), nor for accuracy (χ2 (2) = 4.27, p = 0.12) were found.

For the emotional task, results showed a significant effect of tDCS protocol on reaction time (χ2(2) = 6.15, p = 0.04). The 3mA tDCS was slower than sham (b = -0.014, standard error (SE) = 0.01, p = 0.01), whereas no significant difference was found for the other comparisons (3mA vs. 1.5mA: b = -0.007, SE = 0.01, p = 0.2 and 1.5mA vs. sham: b = -0.007, SE = 0.01, p = 0.21). Accuracy analyses also revealed a significant effect of tDCS protocol (χ2(2) = 12.8, p = 0.001), with the 3mA tDCS presenting higher accuracy compared to 1.5mA tDCS (Odds Ratio (OR) = 0.87, SE = 0.043, p = 0.007) and sham (OR = 0.84, SE = 0.042, p < 0.001). No significant difference in accuracy was found between tDCS 1.5mA and sham (OR = 0.96, SE = 0.047, p = 0.42) (Figure 1).

**Figure 1.**
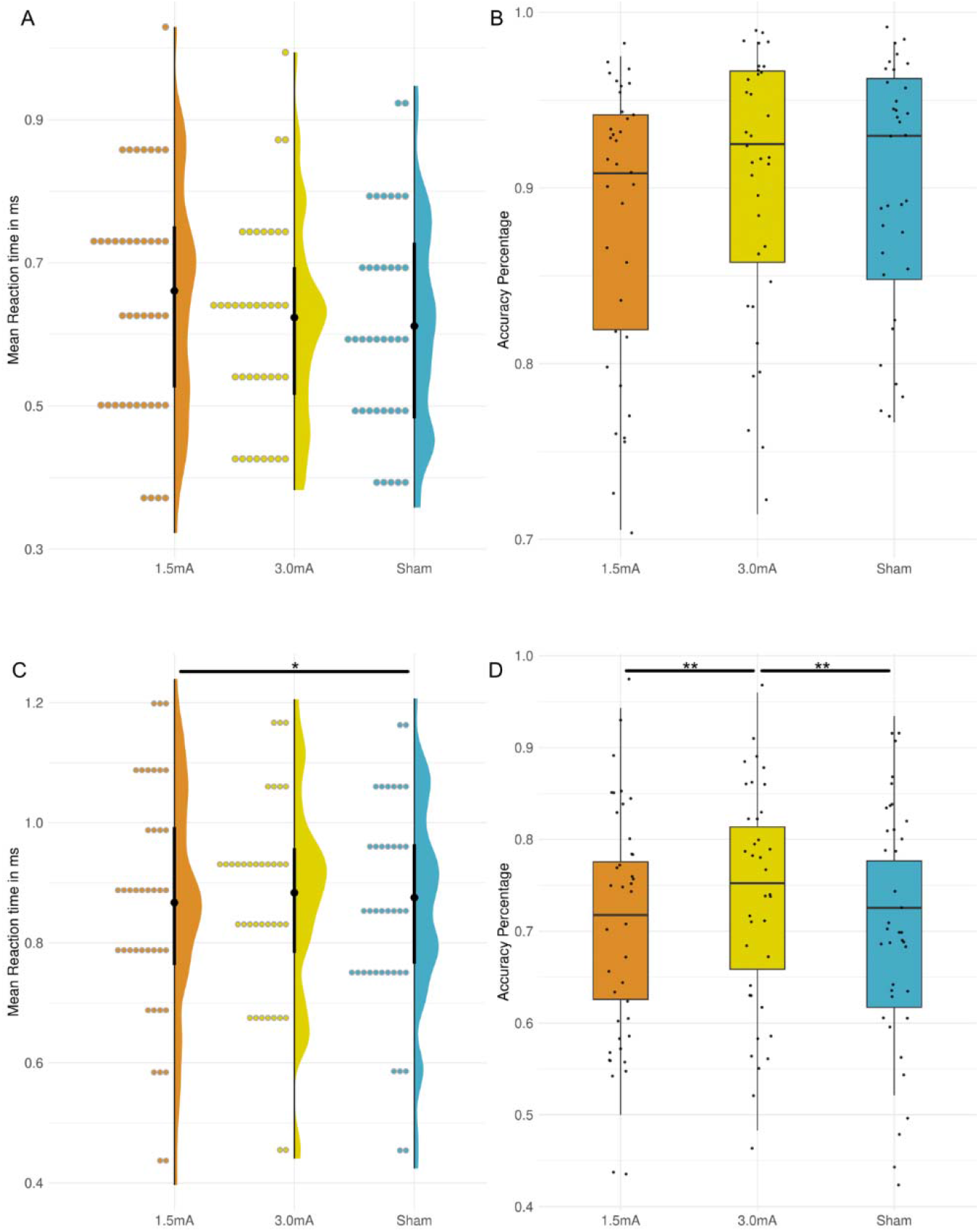
Working memory performance per tDCS protocol. Note: For visualization purposes, the mean reaction time and accuracy per subject and condition were used in Figures 1A 1B, 1C and 1D (whereas statistical analyses were based on the raw data). In the reaction time graph, the median is denoted by the back point, while the interquartile range is represented by the black bars. Similarly, for accuracy data, the median values are represented by the dark line in the box plots, while the interquartile range is represented by the colorful boxes.

A significant effect of ‘session’ was detected for both reaction time (p < 0.001) and accuracy (p < 0.001) in both tasks (Sup. Material - Appendix 7).

### Individual E-field magnitude analysis

The E-field analyses were performed on 37 individuals (one participant was excluded due to erroneous segmentation and two were drop-outs) for both active tDCS protocols, totaling in 74 simulations. Figure 2 shows the individual E-field strength distribution across all 37 heads using 3mA (Sup. material - Appendix 8 shows the E-field distribution of 1.5mA). Participants receiving the higher E-field magnitude in the DLPFC areas with 3mA presented a mean of 0.57 V/m, whereas the lowest E-field magnitude presented a mean value of 0.37V/m (or 0.285V/m and 0.185V/m, respectively, for the 1.5mA protocol). The peak E-field magnitude (99th percentile) of all participants was predominantly situated in the medial part of the DLPFC and is displayed in the Sup. Material - Appendix 8.

**Figure 2.**
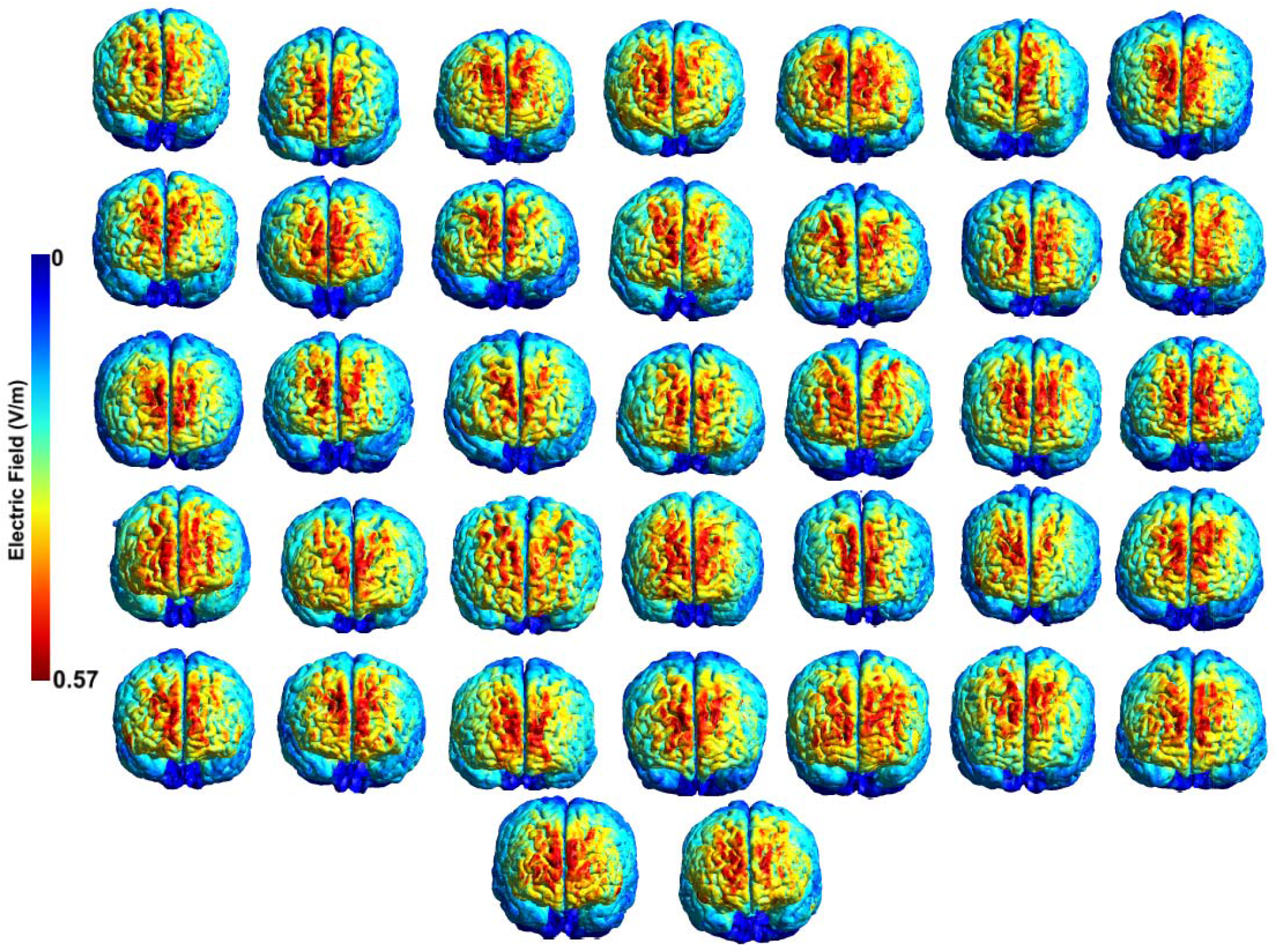
Individual E-field distribution. Note: The presented figure illustrates the distribution of the E-field range for the tDCS protocol at an intensity of 3mA. For the E-field distribution of the protocol at 1.5mA, please see the Supplementary Material - Appendix 8. It should be emphasized that the individual distribution of the E-field remains consistent for both tDCS protocols and the only variation lies in the scaling of the E-field magnitude, given that the E-field intensity for the 3mA tDCS protocol is twice that of the 1.5mA protocol.

### Working memory performance and E-field magnitude

A significant association between reaction time of the active tDCS protocols (minus sham) and E-field magnitude of all brain regions of interest (Table 1; Figure 3) was found after correcting for multiple comparisons, showing that people with higher E-field magnitude in these regions of interest responded faster. For accuracy, no significant association was found (Table 1; Sup. Material - Appendix 9).

**Table 1.**
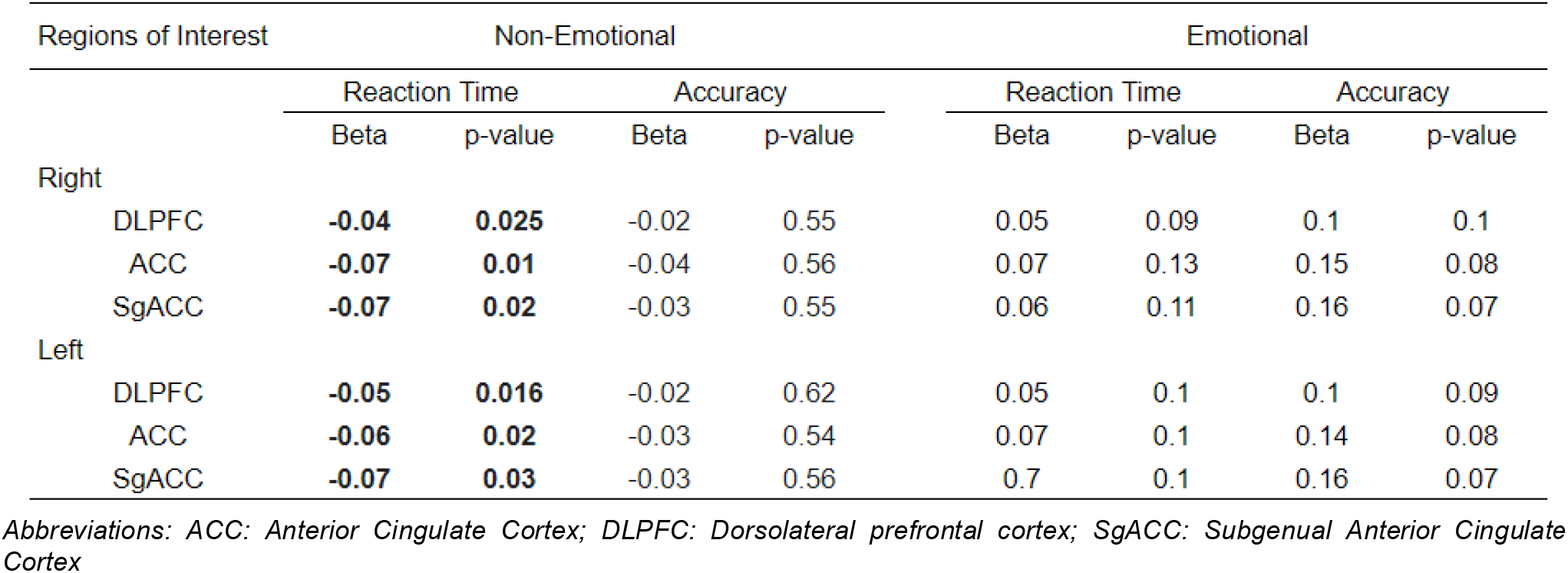
Association between E-field strength in brain regions of interest and working memory performance.

| Regions of Interest |  | Non-Emotional |  |  |  | Emotional |  |  |  |
| --- | --- | --- | --- | --- | --- | --- | --- | --- | --- |
|  |  | Reaction Time |  | Accuracy |  | Reaction Time |  | Accuracy |  |
|  |  | Beta | p-value | Beta | p-value | Beta | p-value | Beta | p-value |
| Right |  |  |  |  |  |  |  |  |  |
|  | DLPFC | **-0.04** | **0.025** | -0.02 | 0.55 | 0.05 | 0.09 | 0.1 | 0.1 |
|  | ACC | **-0.07** | **0.01** | -0.04 | 0.56 | 0.07 | 0.13 | 0.15 | 0.08 |
|  | SgACC | **-0.07** | **0.02** | -0.03 | 0.55 | 0.06 | 0.11 | 0.16 | 0.07 |
| Left |  |  |  |  |  |  |  |  |  |
|  | DLPFC | **-0.05** | **0.016** | -0.02 | 0.62 | 0.05 | 0.1 | 0.1 | 0.09 |
|  | ACC | **-0.06** | **0.02** | -0.03 | 0.54 | 0.07 | 0.1 | 0.14 | 0.08 |
|  | SgACC | **-0.07** | **0.03** | -0.03 | 0.56 | 0.7 | 0.1 | 0.16 | 0.07 |

Abbreviations: ACC: Anterior Cingulate Cortex; DLPFC: Dorsolateral prefrontal cortex; SgACC: Subgenual Anterior Cingulate Cortex

**Figure 3.**
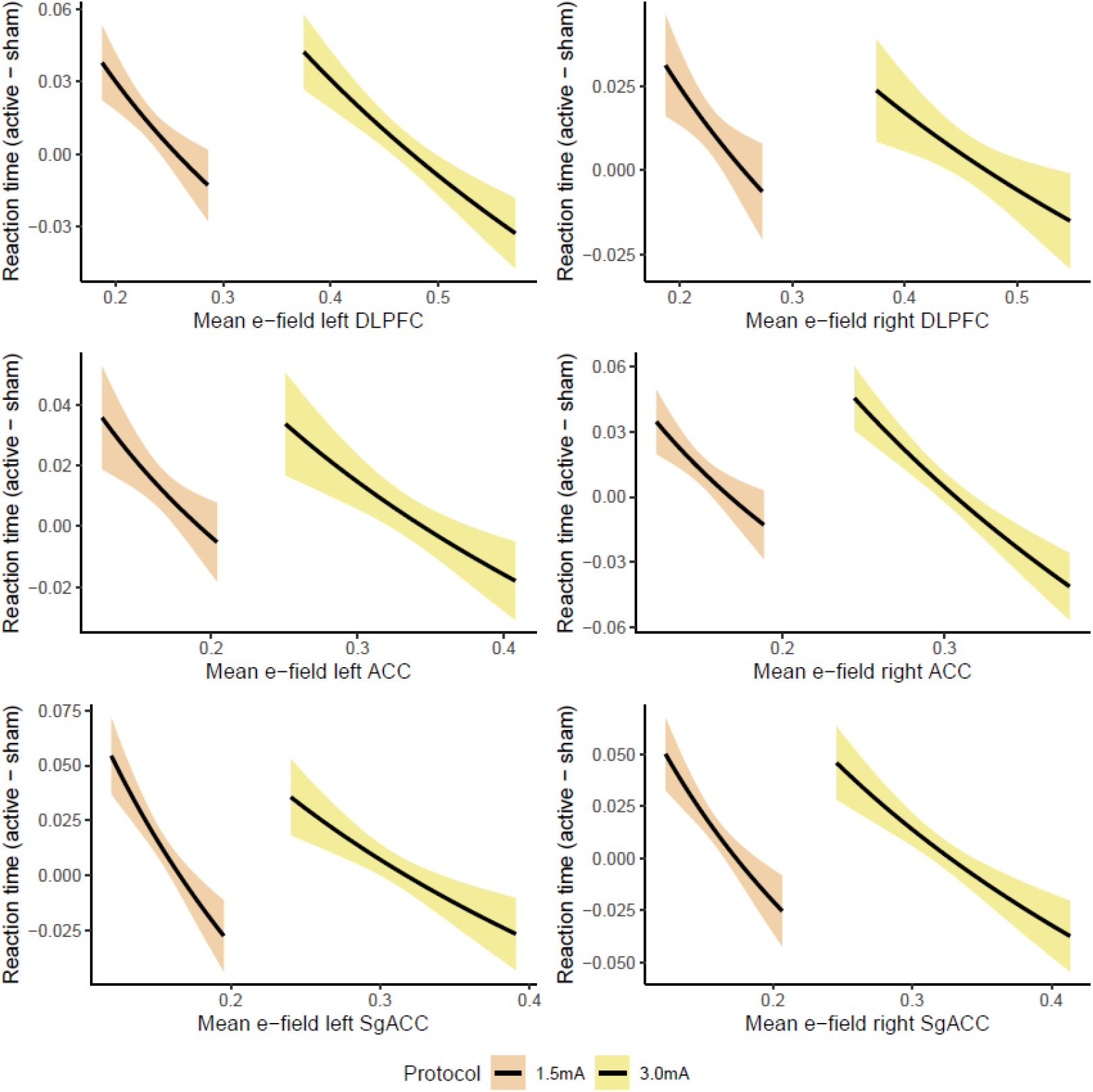
Association between brain regions of interest and reaction time in the non-emotional paradigm

The emotional 3-back results showed no association between reaction time/accuracy and E-field strength in the brain regions investigated (Table 1; Sup. Material - Appendix 9).

### Follow-up analysis: higher vs. lower E-fields

For the non-emotional task, individuals with above-median (*n*=18) E-fields values showed faster reaction time (beta = -0.04, SE = 0.007, p < 0.001) while no difference was found for accuracy (beta = 0.01, SE = 0.01, p = 0.25). For the emotional task, people with above-median E-fields strength in the left DLPFC presented faster reaction time (beta = -0.02, SE = 0.009, *p* = 0.025) and more accurate responses (beta = 0.05, SE = 0.01, p < 0.001) compared to the below-median (n=17) subjects (Figure 4).

**Figure 4.**
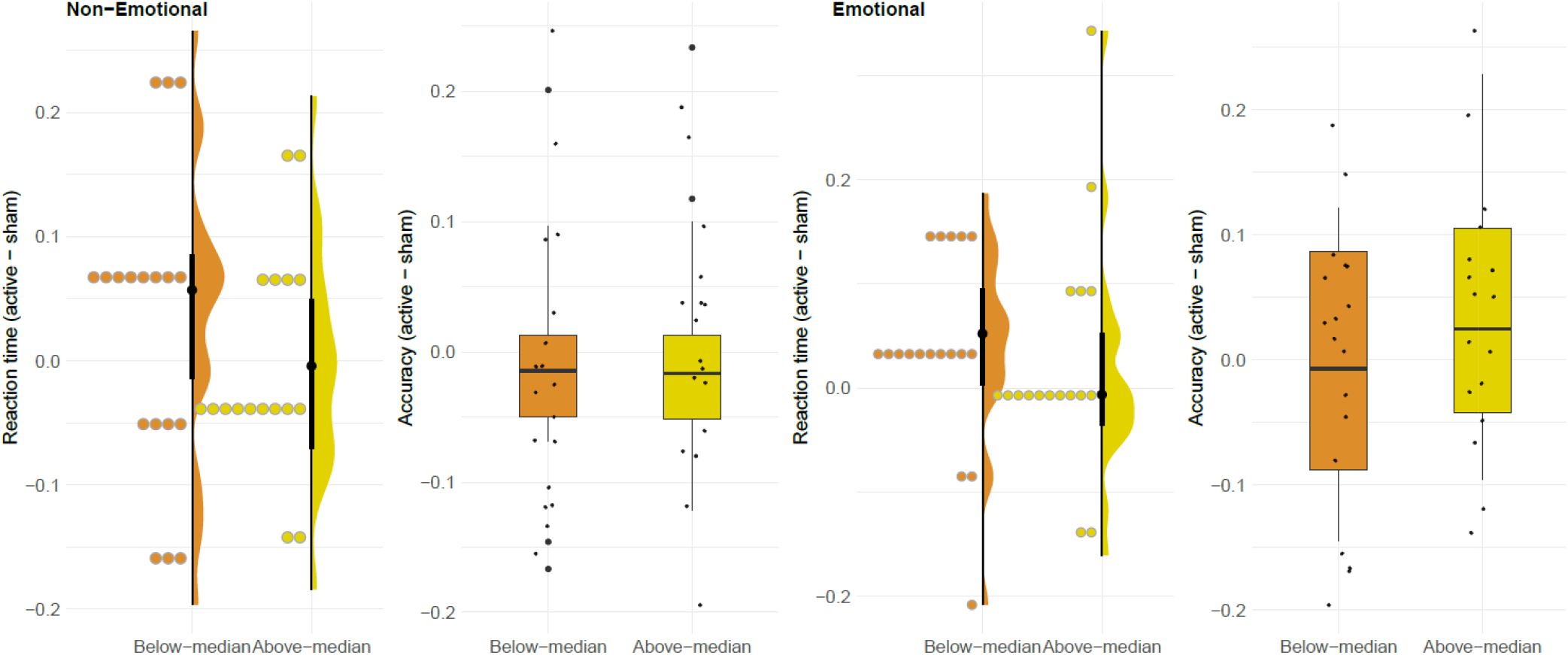
Above and below-median E-field values in the left DLPFC.

### Adverse Effects

No participant presented serious adverse events, such as dizziness or seizure. The 3mA protocol showed higher incidences of local pain, tingling and burning sensation (Sup. Material - Appendix 10).

## Discussion

Here, we investigated whether individual E-field magnitude of distinct active tDCS intensities (1.5mA or 3mA) in brain regions of interest would be associated with working memory performance of healthy subjects across different paradigms. Two statistical approaches were applied aiming to comprehensively overview the effects of prefrontal tDCS on working memory. First, we checked the differences among tDCS protocols only. Results showed that active tDCS protocols did not significantly improve working memory in the non-emotional task, whereas the 3mA modulated performance in the emotional paradigm. In the second approach, the individual E-field magnitude in the brain regions of interest were associated with working memory performance. Results showed that participants with higher E-field magnitude exhibited faster responses in the non-emotional paradigm, whereas no association with the emotional working memory task was found. A follow-up analysis confirmed that individuals with relatively increased E-field in the left DLPFC experienced overall better working memory performance. All in all, the results show that prefrontal tDCS effects on working memory might vary depending on the individual magnitude of E-field in the targeted stimulation area.

Previous studies showed that higher E-fields in brain regions of interest might be associated with more beneficial outcomes in both depressed and healthy samples (Caulfield et al., 2021, 2022b)(Caulfield et al., 2020, 2022b; Suen et al., 2020). Our findings support this prior evidence and add to the growing body of evidence emphasizing the importance of E-field magnitude as a potent source of interindividual variability in the clinical tDCS field. In this sense, the visual correlation between E-fields and reaction time in the non-emotional paradigm (Figure XX) suggests that people receiving higher E-field in the brain regions of interest, independently of the fixed electric current applied, profits more from the intervention. In turn, the 3mA intensity seems to present an increased degree of E-field variability among individuals, which may potentially alleviate the inconsistency observed in cognitive performance, thereby permitting a more consistent outcome.

Our results reinforce the variability between summed scores and individual-level results (i.e., individual E-field magnitude) in the tDCS field (Polanía et al., 2018). Therefore, it suggests that the individual E-field distribution holds potential significance in understanding the overall heterogeneous and in particular null effects of prefrontal tDCS in cognition and possibly also in depression. This is because significant results might be mitigated when considering summed scores as compared to taking into account inter-individual variation in E-field distribution. The findings of the follow-up analysis (i.e. below or above median E-field) reinforce this interpretation and are in line with a recent study which showed similar results with older healthy adults (Caulfield et al., 2022b).

We were unable to demonstrate that participants with higher E-field strength in the ACC and the sgACC presented better performance in the emotional task. The cingulate cortex, specifically the ACC and the sgACC, are directly associated with emotion processes (Rolls, 2019) and present activity dysfunction in depressive episodes (Benschop et al., 2022). A previous study revealed that higher E-fields in the DLPFC and ACC of depressed patients were associated with a decrease in negative affect (Suen et al., 2020). Within this context, we hypothesize that the association between emotional outcomes with the E-field of deeper brain regions might be related to the amount of current that first penetrates the targeted area (i.e., DLPFC). Therefore, stronger E-fields might be needed to achieve deeper areas. The variability in tissue types introduces differences in the penetration of current into the brain and can impact the amount of current flowing into specific brain regions. However, although we have not hypothesized that the the E-field magnitude in the ACC and sgACC would be associated with performance in the non-emotional paradigm, our findings are supported by a previous study showing that the dorsal cingulate cortex is consistently activated during n-back paradigms (Wang et al., 2019).

Overall, our study provides additional evidence that tDCS can effectively modulate brain activity. However, its effects are prone to a source of inter-individual variability that may influence its overall efficacy (Wiethoff et al., 2014)(Agboada et al., 2020; Bastani & Jaberzadeh, 2013)(Jamil et al., 2017), as systematically demonstrated here. Our results can aid in identifying sources of variability that could be mitigated in future clinical investigations. Specifically, a computational modeling approach might assist in future research to prospectively identify the best electric current for each subject and can help to individualize tDCS intensity by reverse E-field calculation in the future.

Moreover, we posit that utilizing more cognitively demanding tasks - such as the emotional 3-back - may reveal more robust effects of tDCS and should be used in future studies to reduce variability. Furthermore, even though we targeted the bilateral DLPFC, this approach might not be the ideal method to stimulate the DLPFC region per se, as the current peak between the anode and cathode relies on the medial part of the DLPFC (see Sup. Material - Appendix 6). Future studies should be encouraged to investigate novel target positions, as proposed in a recent meta-analysis (Wischnewski et al., 2021).

Finally, our study provides evidence that the usage of 3 mA tDCS is a valid strategy to induce E-field magnitudes of ∼ 0.5 V/m, which are often linked to the neural effects of tDCS.

## Limitation

The present E-field analysis only investigated the association between the overall E-field magnitude and working memory performance, but not the differences between currents of 1.5mA and 3mA. Moreover, it is possible that the working memory tasks utilized in this study are susceptible to ceiling effects, which could potentially limit the detection of significant differences across tDCS protocols. Finally, due to the within-subject design, blinding was not assessed as it could increase participants’ awareness regarding the intervention protocol in the subsequent session.

## Conclusion

We systematically evaluated the association between individual magnitude of E-field produced by tDCS (1.5mA, 3mA and sham) in brain regions of interest (DLPFC, ACC, sgACC) and working memory performance of two different paradigms (emotional and non-emotional). Our study confirms that prefrontal tDCS effects in healthy individuals modulates working memory processes, but emphasizes the importance of E-field magnitude as an essential physical agent underlying the interindividual variability in its effectiveness. Therefore, these results suggest that individual E-field distribution holds potential significance in understanding the overall heterogeneity of prefrontal tDCS in cognition and also in depression effects, which might be improved by personalizing current intensity according to head anatomy, although this hypothesis should be confirmed by further studies.

## Acknowledgment

We thank our Masters students Mathilde Nay and Xander Cornelis for collecting and organizing the data of this work.

## Disclosure

ARB receives grants from the National Council for Scientific and Technological Development (PQ-1B), and FAPESP (Grants: 2018/10861-7, 2019/06009-6). ARB has a small equity of Flow^™^, whose devices were not used in the present study. The LIM-27 laboratory receives grants from the Associação Beneficente Alzira Denise Hertzog da Silva. LBR is currently supported by a Research Foundation Flanders (FWO) grant (G0F4619N). MAV receives funding from the FWO and from Ghent University (Grants: G0F4619N and BOF17/STA/030, respectively). SDS is funded by a FWO-Flanders PhD fellowship (Grant Number: 11J7521N). MSL is funded by FAPESP (grant number: 2021/10574-0). The other authors have no conflicts of interest to disclose.

## Notes

### Competing Interest Statement

The authors have declared no competing interest.

## References

Agboada, D., Mosayebi-Samani, M., Kuo, M.-F., & Nitsche, M. A. (2020). Induction of long-term potentiation-like plasticity in the primary motor cortex with repeated anodal transcranial direct current stimulation -Better effects with intensified protocols? Brain Stimulation, 13(4), 987–997.

Albizu, A., Fang, R., Indahlastari, A., O’Shea, A., Stolte, S. E., See, K. B., Boutzoukas, E. M., Kraft, J. N., Nissim, N. R., & Woods, A. J. (2020). Machine learning and individual variability in electric field characteristics predict tDCS treatment response. Brain Stimulation, 13(6), 1753–1764.

Aparício, L. V. M., Guarienti, F., Razza, L. B., Carvalho, A. F., Fregni, F., & Brunoni, A. R. (2016). A Systematic Review on the Acceptability and Tolerability of Transcranial Direct Current Stimulation Treatment in Neuropsychiatry Trials. Brain Stimulation, 9(5), 671–681.

Bastani, A., & Jaberzadeh, S. (2013). a-tDCS differential modulation of corticospinal excitability: the effects of electrode size. Brain Stimulation, 6(6), 932–937.

Beck, A. T., Ward, C. H., Mendelson, M., Mock, J., & Erbaugh, J. (1961). An inventory for measuring depression. Archives of General Psychiatry, 4, 561–571.

Benschop, L., Vanhollebeke, G., Li, J., Leahy, R. M., Vanderhasselt, M.-A., & Baeken, C. (2022). Reduced subgenual cingulate-dorsolateral prefrontal connectivity as an electrophysiological marker for depression. Scientific Reports, 12(1), 16903.

Blumberger, D. M., Vila-Rodriguez, F., Thorpe, K. E., Feffer, K., Noda, Y., Giacobbe, P., Knyahnytska, Y., Kennedy, S. H., Lam, R. W., Daskalakis, Z. J., & Downar, J. (2018). Effectiveness of theta burst versus high-frequency repetitive transcranial magnetic stimulation in patients with depression (THREE-D): a randomised non-inferiority trial. The Lancet, 391(10131), 1683–1692.

Bulubas, L., Padberg, F., Bueno, P. V., Duran, F., Busatto, G., Amaro, E., Jr, Benseñor, I. M., Lotufo, P. A., Goerigk, S., Gattaz, W., Keeser, D., & Brunoni, A. R. (2019). Antidepressant effects of tDCS are associated with prefrontal gray matter volumes at baseline: Evidence from the ELECT-TDCS trial. Brain Stimulation, 12(5), 1197–1204.

Caulfield, K. A., Badran, B. W., DeVries, W. H., Summers, P. M., Kofmehl, E., Li, X., Borckardt, J. J., Bikson, M., & George, M. S. (2020). Transcranial electrical stimulation motor threshold can estimate individualized tDCS dosage from reverse-calculation electric-field modeling. Brain Stimulation, 13(4), 961–969.

Caulfield, K. A., Indahlastari, A., Nissim, N. R., Lopez, J. W., Fleischmann, H. H., Woods, A. J., & George, M. S. (2022a). Electric Field Strength From Prefrontal Transcranial Direct Current Stimulation Determines Degree of Working Memory Response: A Potential Application of Reverse-Calculation Modeling? Neuromodulation: Journal of the International Neuromodulation Society, 25(4), 578–587.

Caulfield, K. A., Indahlastari, A., Nissim, N. R., Lopez, J. W., Fleischmann, H. H., Woods, A. J., & George, M. S. (2022b). Electric Field Strength From Prefrontal Transcranial Direct Current Stimulation Determines Degree of Working Memory Response: A Potential Application of Reverse-Calculation Modeling? Neuromodulation: Journal of the International Neuromodulation Society, 25(4), 578–587.

Caulfield, K. A., Li, X., & George, M. S. (2021). A reexamination of motor and prefrontal TMS in tobacco use disorder: Time for personalized dosing based on electric field modeling? Clinical Neurophysiology: Official Journal of the International Federation of Clinical Neurophysiology, 132(9), 2199–2207.

Dedoncker, J., Brunoni, A. R., Baeken, C., & Vanderhasselt, M.-A. (2016). A Systematic Review and Meta-Analysis of the Effects of Transcranial Direct Current Stimulation (tDCS) Over the Dorsolateral Prefrontal Cortex in Healthy and Neuropsychiatric Samples: Influence of Stimulation Parameters. Brain Stimulation, 9(4), 501–517.

De Smet, S., Nikolin, S., Moffa, A., Suen, P., Vanderhasselt, M.-A., Brunoni, A. R., & Razza, L. B. (2021). Determinants of sham response in tDCS depression trials: a systematic review and meta-analysis. Progress in Neuro-Psychopharmacology &Biological Psychiatry, 109, 110261.

Etkin, A., Egner, T., & Kalisch, R. (2011). Emotional processing in anterior cingulate and medial prefrontal cortex. Trends in Cognitive Sciences, 15(2), 85–93.

Fan, L., Li, H., Zhuo, J., Zhang, Y., Wang, J., Chen, L., Yang, Z., Chu, C., Xie, S., Laird, A. R., Fox, P. T., Eickhoff, S. B., Yu, C., & Jiang, T. (2016). The Human Brainnetome Atlas: A New Brain Atlas Based on Connectional Architecture. Cerebral Cortex, 26(8), 3508–3526.

Fox, M. D., Buckner, R. L., White, M. P., Greicius, M. D., & Pascual-Leone, A. (2012). Efficacy of transcranial magnetic stimulation targets for depression is related to intrinsic functional connectivity with the subgenual cingulate. Biological Psychiatry, 72(7), 595–603.

Indahlastari, A., Albizu, A., Kraft, J. N., O’Shea, A., Nissim, N. R., Dunn, A. L., Carballo, D., Gordon, M. P., Taank, S., Kahn, A. T., Hernandez, C., Zucker, W. M., & Woods, A. J. (2021). Individualized tDCS modeling predicts functional connectivity changes within the working memory network in older adults. Brain Stimulation, 14(5), 1205–1215.

Jamil, A., Batsikadze, G., Kuo, H.-I., Labruna, L., Hasan, A., Paulus, W., & Nitsche, M. A. (2017). Systematic evaluation of the impact of stimulation intensity on neuroplastic after-effects induced by transcranial direct current stimulation. The Journal of Physiology, 595(4), 1273–1288.

Koenigs, M., & Grafman, J. (2009). The functional neuroanatomy of depression: distinct roles for ventromedial and dorsolateral prefrontal cortex. Behavioural Brain Research, 201(2), 239–243.

Majdi, A., van Boekholdt, L., Sadigh-Eteghad, S., &Mc Laughlin M. (2022). A systematic review and meta-analysis of transcranial direct-current stimulation effects on cognitive function in patients with Alzheimer’s disease. Molecular Psychiatry, 27(4), 2000–2009.

Miró-Padilla, A., Bueichekú, E., Ventura-Campos, N., Flores-Compañ, M.-J., Parcet, M. A., &Ávila, C. (2019). Long-term brain effects of N-back training: an fMRI study. Brain Imaging and Behavior, 13(4), 1115–1127.

Moors, A., De Houwer, J., Hermans, D., Wanmaker, S., van Schie, K., Van Harmelen, A.-L., De Schryver, M., De Winne, J., & Brysbaert, M. (2013). Norms of valence, arousal, dominance, and age of acquisition for 4,300 Dutch words. Behavior Research Methods, 45(1), 169–177.

Pe, M. L., Raes, F., & Kuppens, P. (2013). The cognitive building blocks of emotion regulation: ability to update working memory moderates the efficacy of rumination and reappraisal on emotion. PloS One, 8(7), e69071.

Pinninti, N. R., Madison, H., Musser, E., & Rissmiller, D. (2003). MINI International Neuropsychiatric Schedule: clinical utility and patient acceptance. European Psychiatry: The Journal of the Association of European Psychiatrists, 18(7), 361–364.

Polanía, R., Nitsche, M. A., & Ruff, C. C. (2018). Studying and modifying brain function with non-invasive brain stimulation. Nature Neuroscience, 21(2), 174–187.

Puonti, O., Van Leemput, K., Saturnino, G. B., Siebner, H. R., Madsen, K. H., & Thielscher, A. (2020). Accurate and robust whole-head segmentation from magnetic resonance images for individualized head modeling. NeuroImage, 219, 117044.

Razza, L. B., Palumbo, P., Moffa, A. H., Carvalho, A. F., Solmi, M., Loo, C. K., & Brunoni, A. R. (2020). A systematic review and meta-analysis on the effects of transcranial direct current stimulation in depressive episodes. Depression and Anxiety, 37(7), 594–608.

Rolls, E. T. (2019). The cingulate cortex and limbic systems for emotion, action, and memory. Brain Structure &Function, 224(9), 3001–3018.

Sallet, J., Mars, R. B., Noonan, M. P., Neubert, F.-X., Jbabdi, S., O’Reilly, J. X., Filippini, N., Thomas, A. G., & Rushworth, M. F. (2013). The organization of dorsal frontal cortex in humans and macaques. The Journal of Neuroscience: The Official Journal of the Society for Neuroscience, 33(30), 12255–12274.

Saturnino, G. B., Madsen, K. H., & Thielscher, A. (2019). Electric field simulations for transcranial brain stimulation using FEM: an efficient implementation and error analysis. Journal of Neural Engineering, 16(6), 066032.

Spielberger, C. D., Gorsuch, R. L., & Lushene, R. E. (1970). STAI Manual for the State-trait Anxiety Inventory (“Self-evaluation Questionnaire”).

Spielberg, J. M., Stewart, J. L., Levin, R. L., Miller, G. A., & Heller, W. (2008). Prefrontal Cortex, Emotion, and Approach/Withdrawal Motivation. Social and Personality Psychology Compass, 2(1), 135–153.

Suen, P. J. C., Doll, S., Batistuzzo, M. C., Busatto, G., Razza, L. B., Padberg, F., Mezger, E., Bulubas, L., Keeser, D., Deng, Z.-D., & Brunoni, A. R. (2020). Association between tDCS computational modeling and clinical outcomes in depression: data from the ELECT-TDCS trial. European Archives of Psychiatry and Clinical Neuroscience. https://doi.org/10.1007/s00406-020-01127-w

Van Hoornweder, S., Nuyts, M., Frieske, J., Verstraelen, S., Meesen, R. L. J., & Caulfield, K. A. (2023). A Systematic Review and Large-Scale tES and TMS Electric Field Modeling Study Reveals How Outcome Measure Selection Alters Results in a Person- and Montage-Specific Manner. bioRxiv : The Preprint Server for Biology. https://doi.org/10.1101/2023.02.22.529540

Wang, H., He, W., Wu, J., Zhang, J., Jin, Z., & Li, L. (2019). A coordinate-based meta-analysis of the n-back working memory paradigm using activation likelihood estimation. Brain and Cognition, 132, 1–12.

Wiethoff, S., Hamada, M., & Rothwell, J. C. (2014). Variability in response to transcranial direct current stimulation of the motor cortex. Brain Stimulation, 7(3), 468–475.

Wischnewski, M., Mantell, K. E., & Opitz, A. (2021). Identifying regions in prefrontal cortex related to working memory improvement: A novel meta-analytic method using electric field modeling. Neuroscience and Biobehavioral Reviews, 130, 147–161.

Wörsching, J., Padberg, F., Helbich, K., Hasan, A., Koch, L., Goerigk, S., Stoecklein, S., Ertl-Wagner, B., & Keeser, D. (2017). Test-retest reliability of prefrontal transcranial Direct Current Stimulation (tDCS) effects on functional MRI connectivity in healthy subjects. NeuroImage, 155, 187–201.

Zhang, X., Zhang, R., Lv, L., Qi, X., Shi, J., & Xie, S. (2022). Correlation between cognitive deficits and dorsolateral prefrontal cortex functional connectivity in first-episode depression. Journal of Affective Disorders, 312, 152–158.

