## Supplementary material for "The effects of prefrontal tDCS on working memory associate with the magnitude of the individual electric field in the brain": Supplementary material.docx

**Appendix 1**

**Figure 1.** Study Design

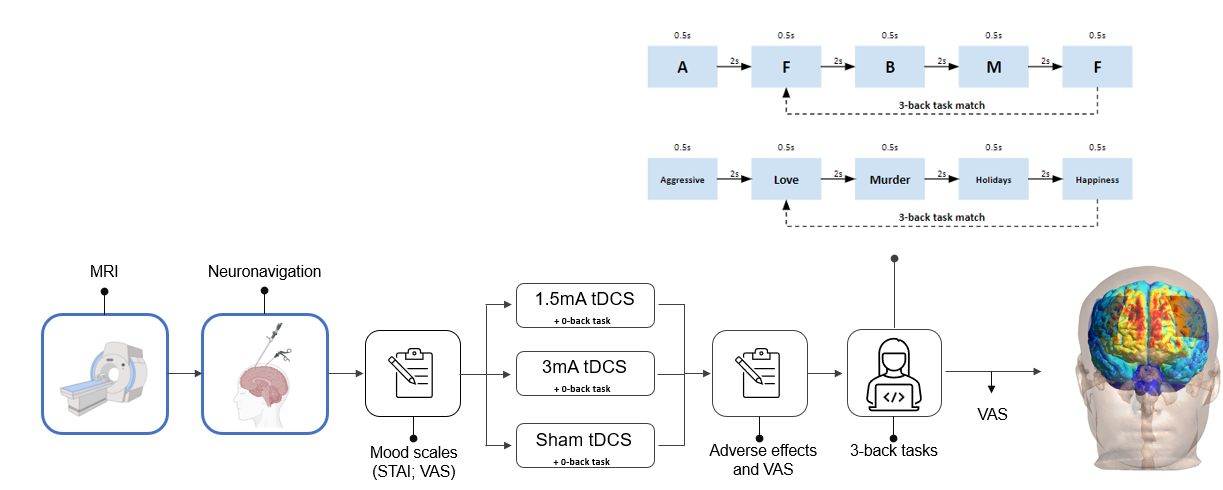

Note: The word stimuli for the emotional 3-back task are here presented in English for better understandment, but they were originally presented in Dutch.

Abbreviation: MRI: Magnetic Resonance Imaging; STAI: State-Trait-ANxiety Inventory; VAS: Visual Analogue Scale.

**Appendix 2**

**Dorsolateral prefrontal cortex (DLPFC)**

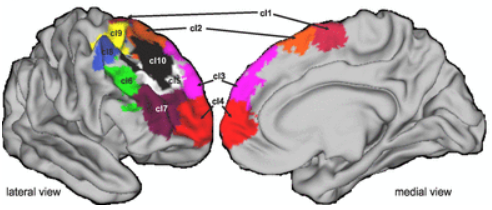

For the DLPFC, we extracted electric field strength of all ten subregions specified in the figure (Sallet et al., 2013) and calculated the mean per subject.

**Anterior Cingulate Cortex (ACC) and Subgenual Anterior Cingulate cortex (SgACC)**

A)	B)
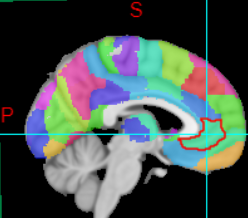

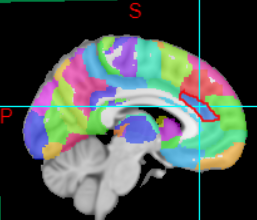

Figure A shows the pregenual ACC (A32p) portion of the Brainnetome atlas (xxx) which here we considered the ACC region. While figure B shows the subgenual ACC (A32sg) portion of the Brainnetome atlas.

**Appendix 3**

Histogram distribution for reaction time of both non-emotional and emotional N-back paradigms:

Non-emotional	Emotional

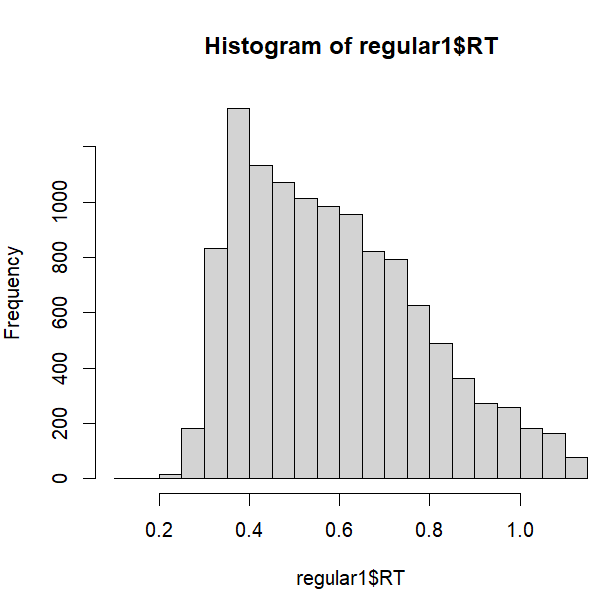

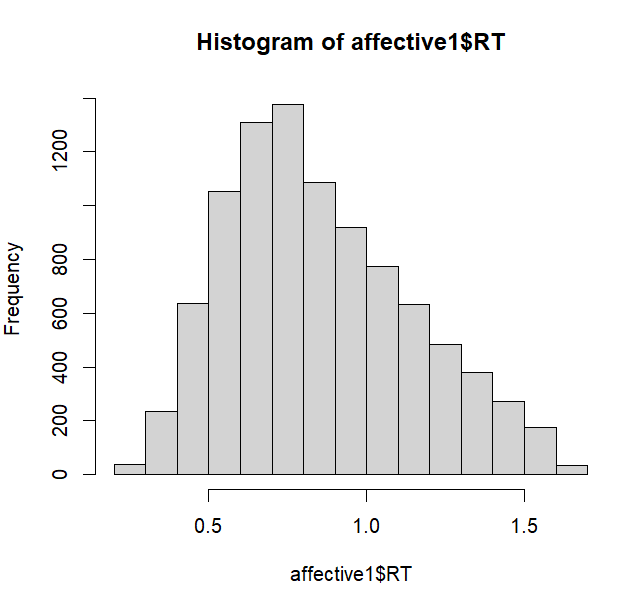

**Working memory performance of Non-emotional paradigm per tDCS protocol:**

Reaction time

fit1<- glmer(RT ~ Protocol + session + (1 | Subject),data=non-emotional, family = Gamma(link = "identity")
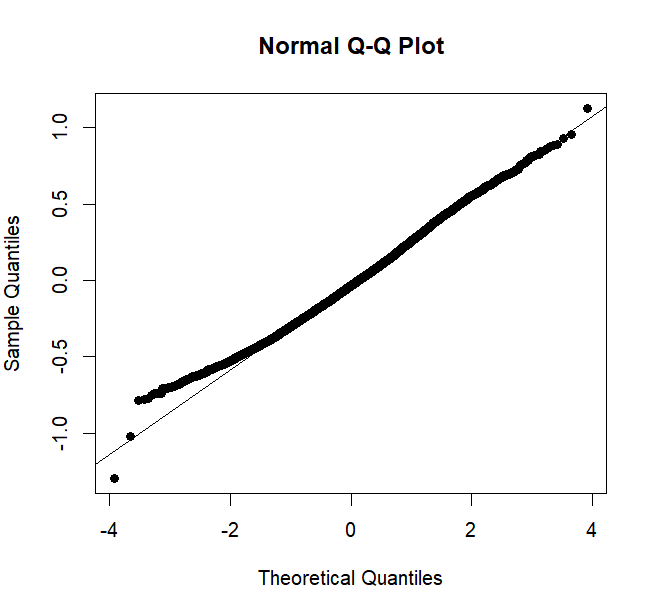

Anova (fit1)

Accuracy

fit2<- glmer(acc ~ Protocol + session + (1|Subject), data = non-emotional, family=binomial())

Anova (fit2)

Both ‘Protocol’ and ‘session’ have three levels.

**Working memory performance of Emotional paradigm per tDCS protocol:**

Reaction time

fit1<- glmer(RT ~ Protocol + session + (1 | Subject),data=emotional, family = Gamma(link = "identity")

Anova (fit1)
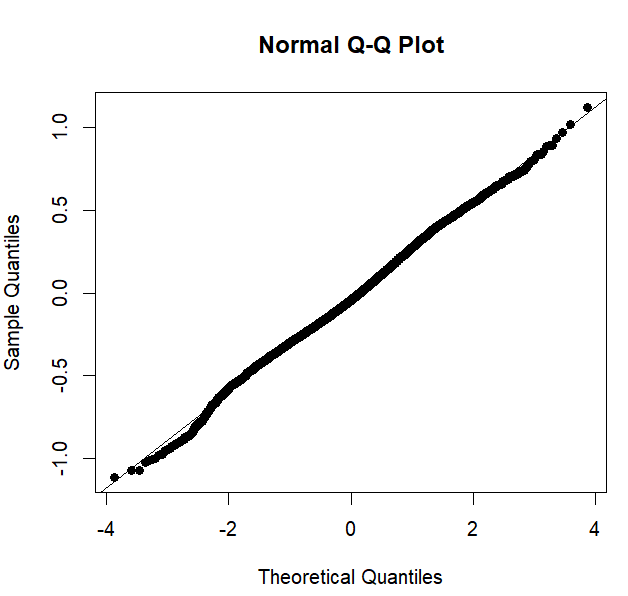

Accuracy

fit2<- glmer(acc ~ Protocol + session + (1|Subject), data = emotional, family=binomial())

Anova (fit2)

Both ‘Protocol’ and ‘session’ have three levels.

**Appendix 4**

**Association between working memory performance and individual E-field:**

For the association between working memory performance and E-field of brain regions of interest, we used linear mixed models (LMM) analysis. For each N-back task there were six LMMs per outcome (reaction time or accuracy), with a total of 12 models per N-back task. Multiple comparison corrections, using the false ratio discovery was therefore performed to avoid false positive findings. LMMs, instead of GLMM were used due to the normal distribution of the data:

Reaction time - Non emotional	Reaction time - Emotional

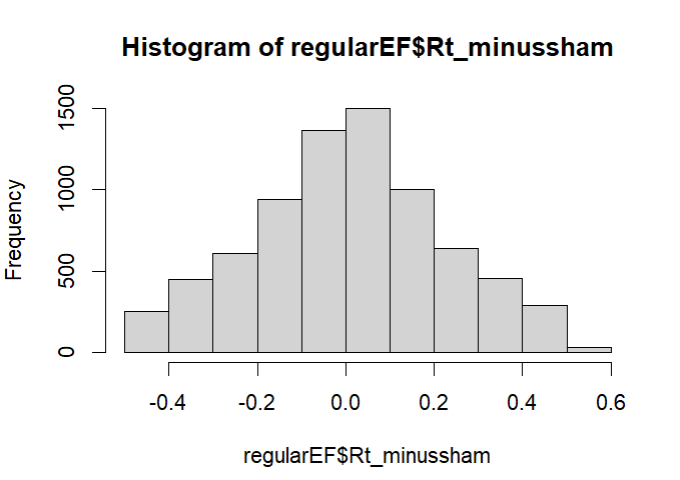

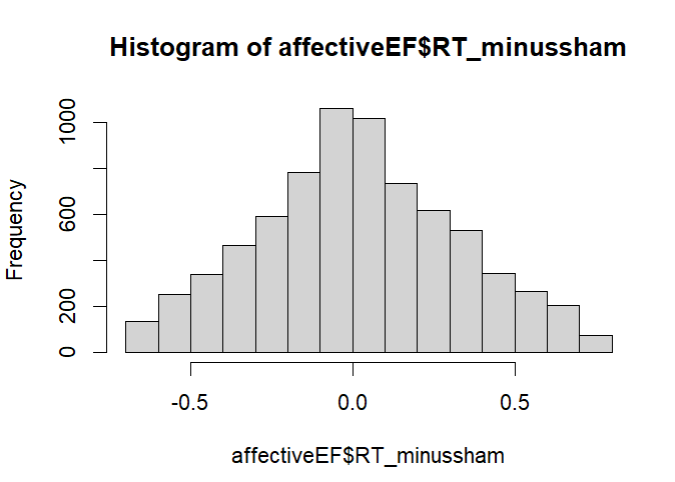

Accuracy - Non emotional	Accuracy - Emotional

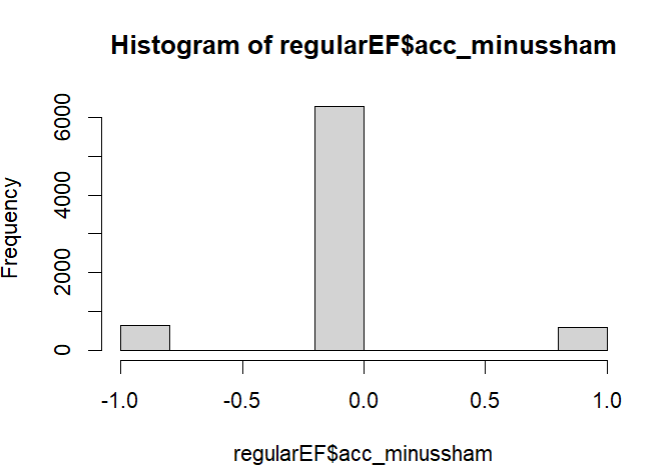

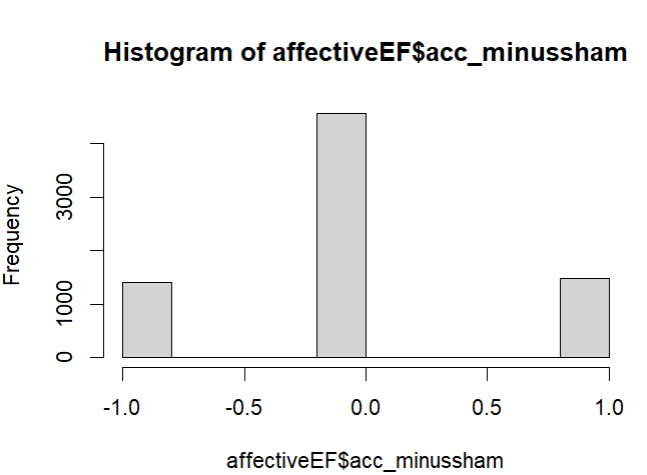

The LMMs applied in our analysis have the following structure:

lm1<-lmer(Rt_minussham~Efieldbrain_region + (1|Subject), data= data)

Anova(lm1)

lm2<-lmer(acc_minussham~Efieldbrain_region + (1|Subject), data= data)

Anova(lm2)

**Association between median and working memory performance:**

Median <- lm(RT_minussham ~ median, data=data)

Anova(Median1)

Median2 <- lm(acc_minussham ~ median, data=data)

Anova(Median2)

Median has two levels (0=below median; 1=above median)

**Tolerability analysis:**

For the tolerability analysis, we used LMMs to fit each adverse effect measured, resulting in a total of 14 models. The adverse effect was the dependent variable, ‘protocol’ was the independent variable, ‘session’ was a fixed factor and ‘subject’ was a random intercept. The models were as following:

fit <- lmer(headache ~ Protocol + Session + (1|Participant),data = data

**Appendix 5**

**Mood effects STAI and VAS**

The mean STAI measure indicated no significant baseline mood differences among protocols (total sham: 44.2; total 1.5mA: 43.9; total 3mA: 44.2, *p*-value = 0.83). For the VAS measure, no significant effect of time ((*F*(2, 295) = 2.85, *p* = 0.07), nor for the interaction between time and tDCS protocol (*F*(4, 296)= 0.19, *p* = 0.094) were found

Mood effects (VAS) per time, per protocol:

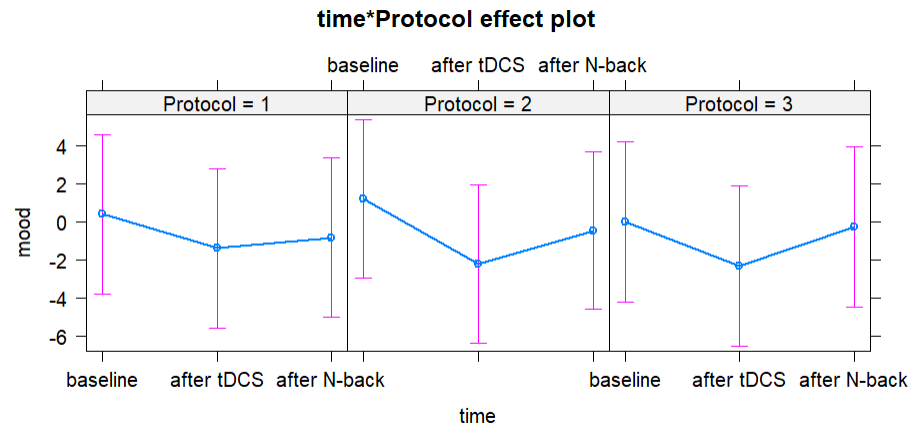

Mood effects (VAS) across session

A significant effect of session was observed (*F*(2, 296)= 6.8, *p* = 0.001), showing that people presented less negative affect across time.

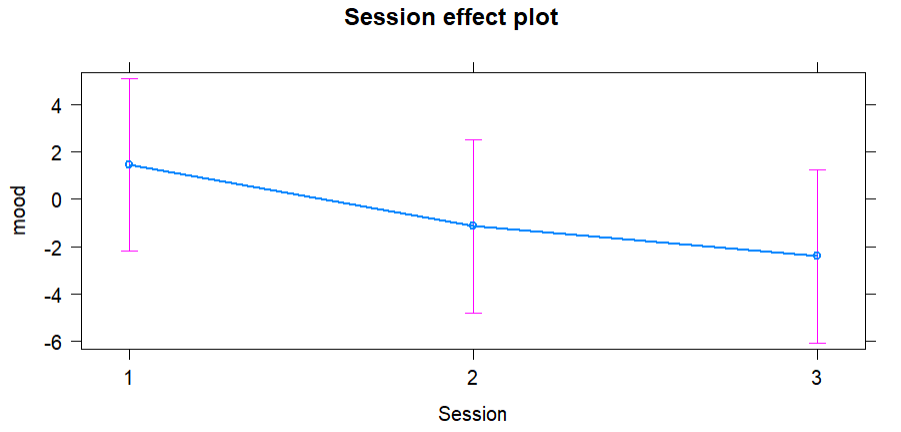

**Appendix 5**

**Table 1.** Mean performance per tDCS protocol in both non-emotional and emotional working memory paradigms.

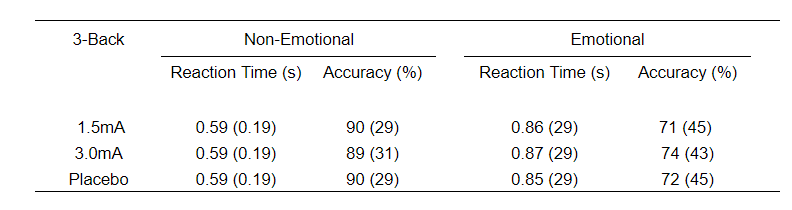

**Appendix 6**

Effect of session in both reaction time and accuracy measure of the non-emotional N-back paradigm.

Reaction time non-emotional	Accuracy non-emotional

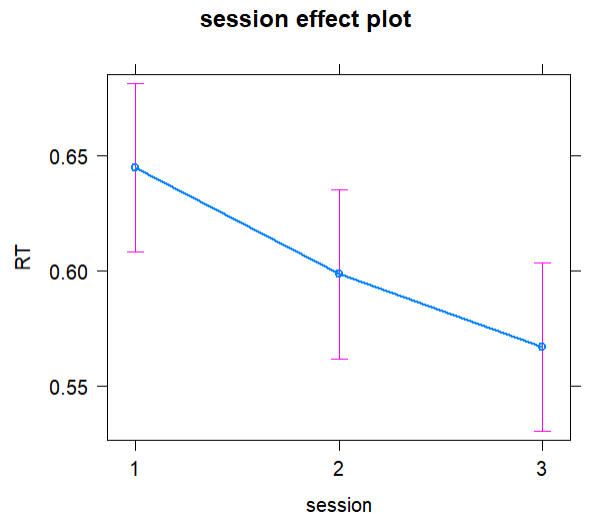

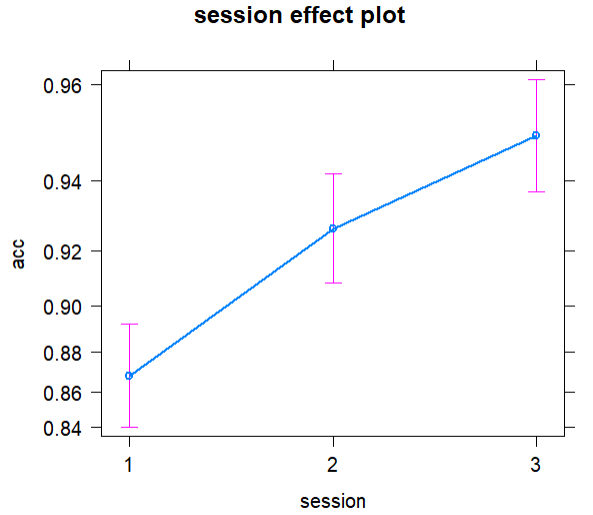

Effect of session in both reaction time and accuracy measure of the emotional N-back paradigm.

Reaction time emotional	Accuracy emotional
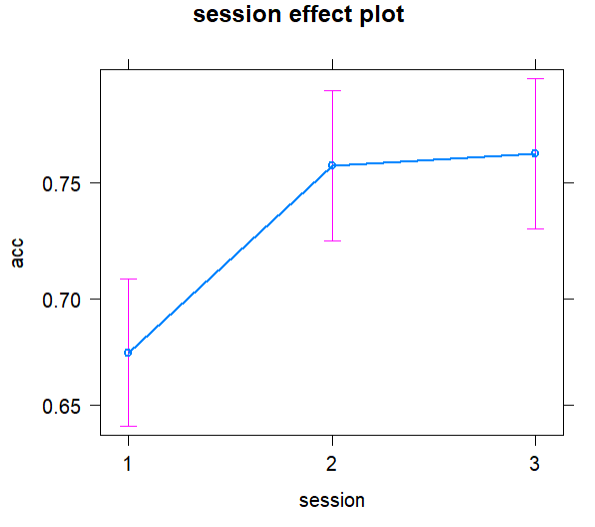

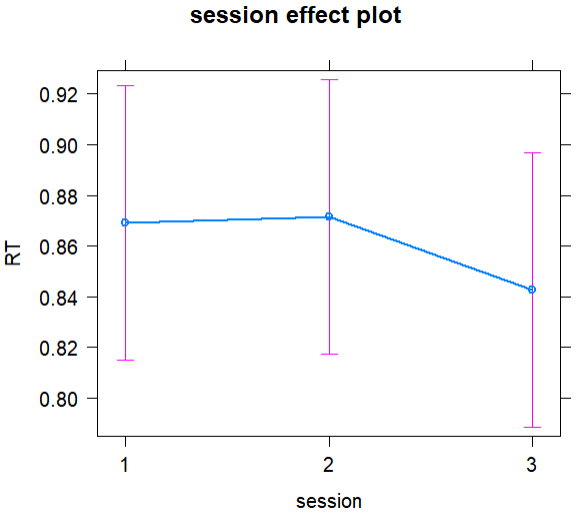

**Appendix 8**

**Figure 1.** Individual distribution of E-field magnitude based on the 1.5mA tDCS protocol.

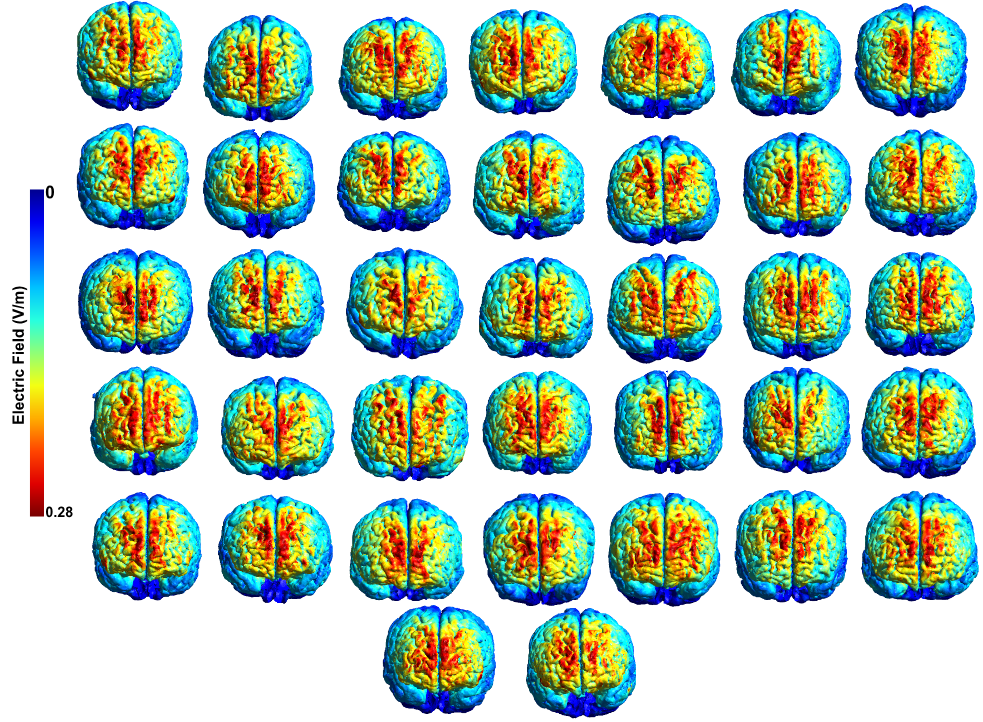

**Figure 2.** Peak E-field magnitude (99 percentile) of all subjects included in our analysis.

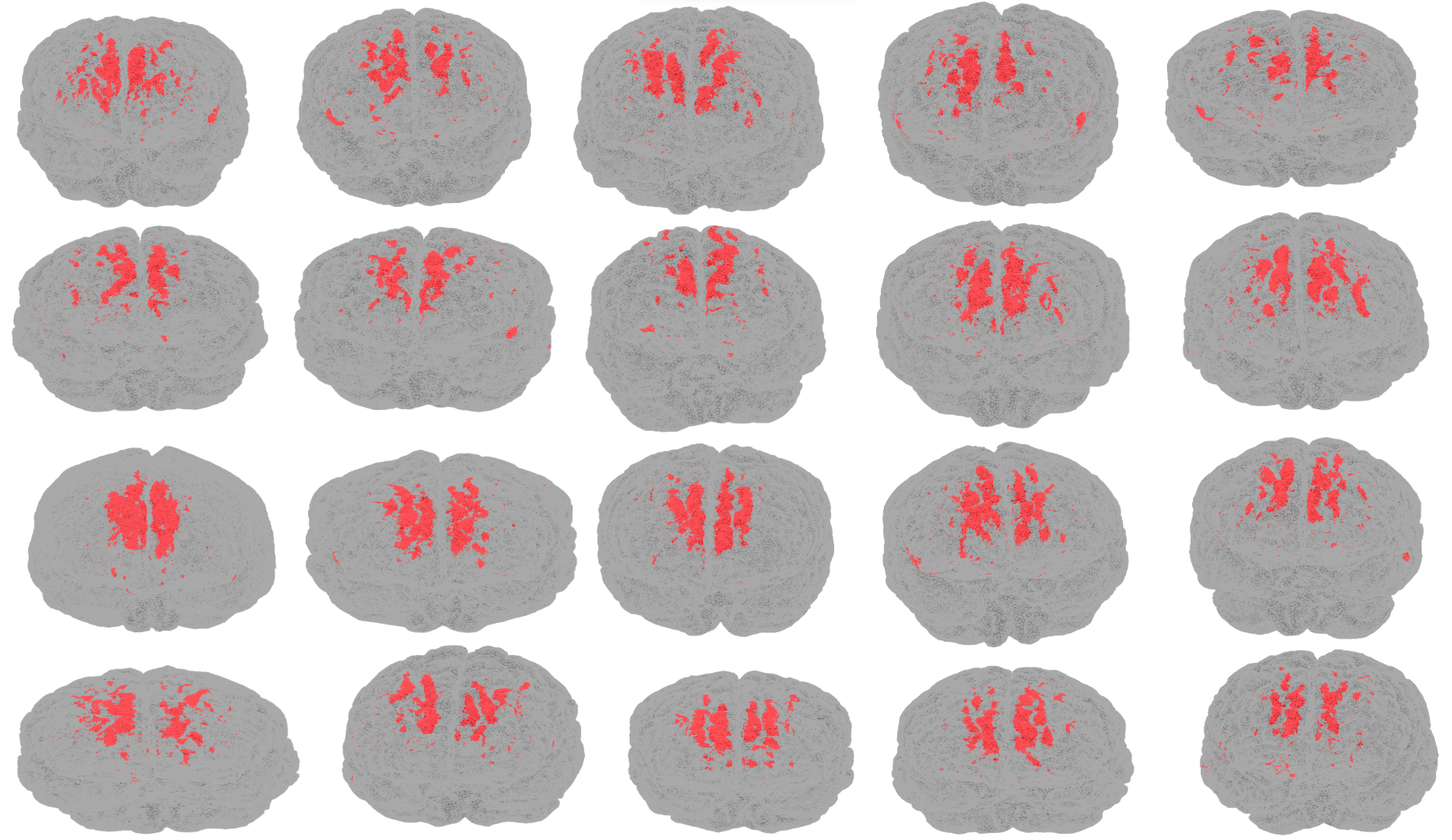

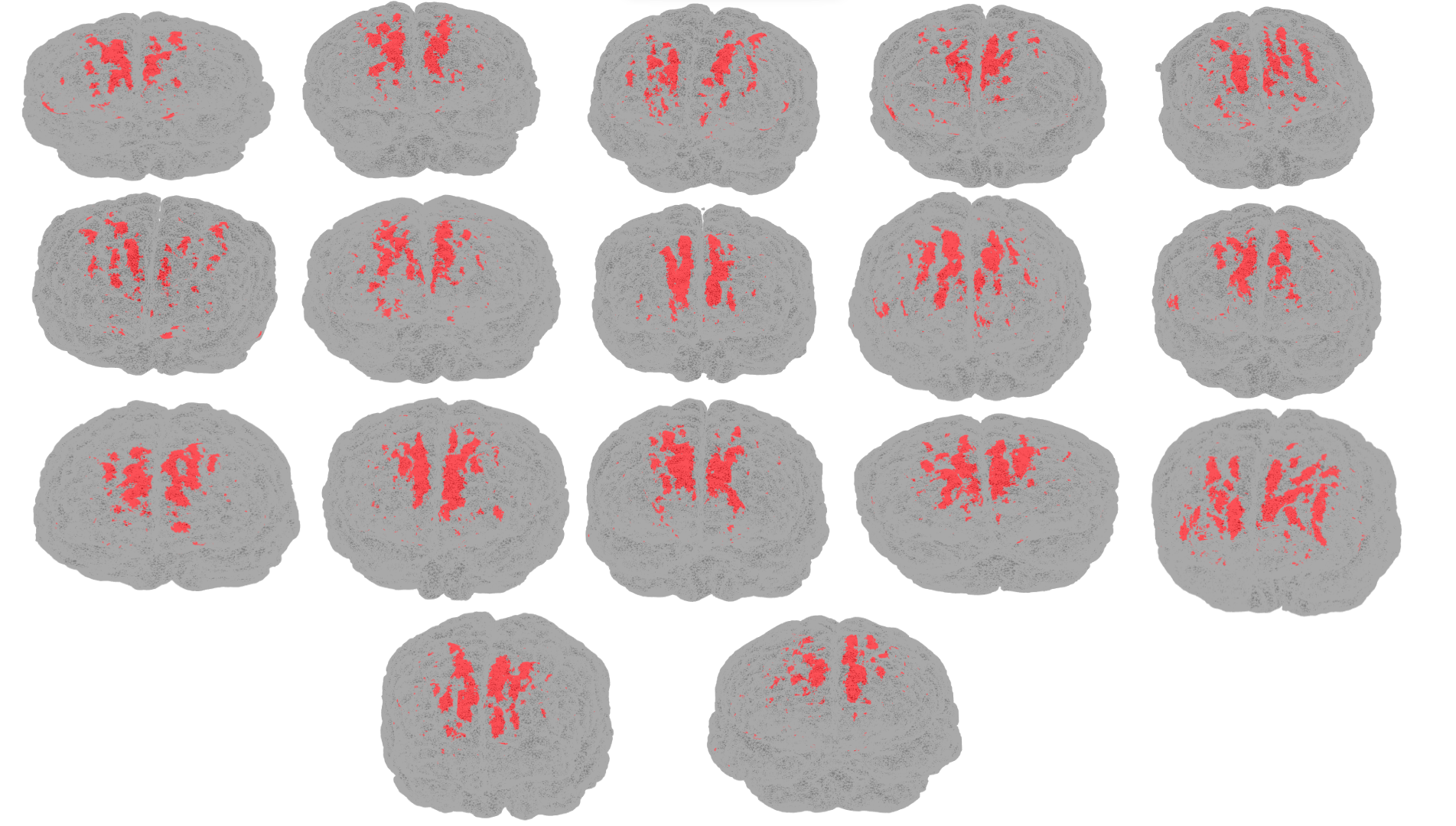

**Appendix 9**

Visualization of the association between E-field in brain regions of interest and accuracy (active minus sham) in the non-emotional 3-task.

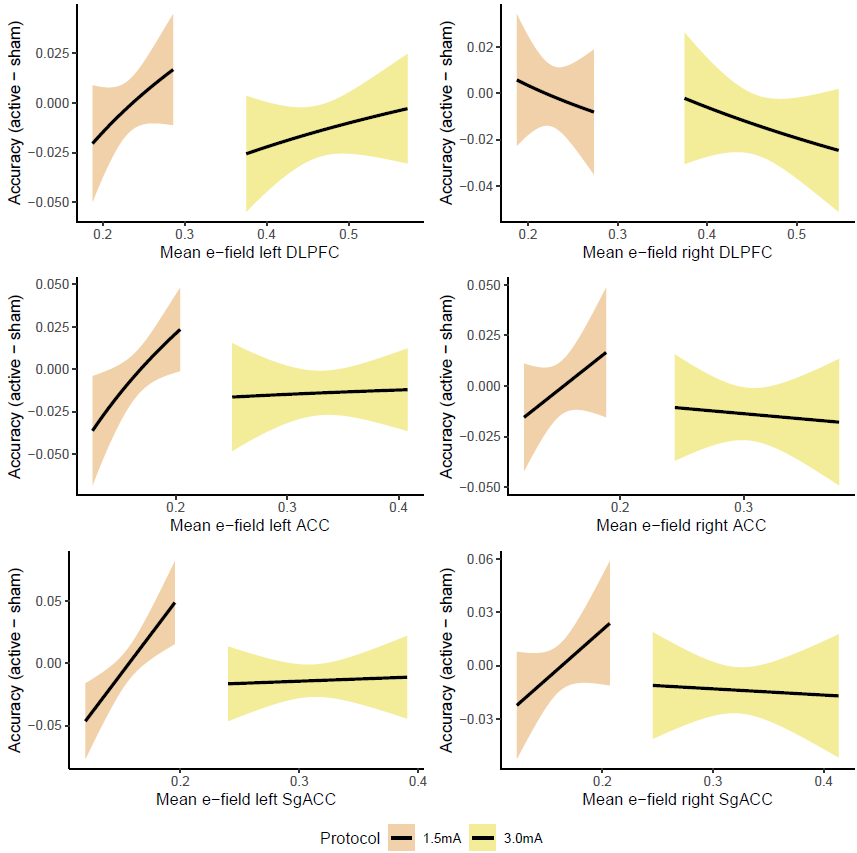

Abbreviation: ACC: Anterior Cingulate Cortex; DLPFC: Dorsolateral Prefrontal Cortex; SgACC: Subgenual Anterior Cingulate Cortex.

Visualization of the association between E-field in brain regions of interest and reaction time performance (active minus sham) in the emotional 3-task.

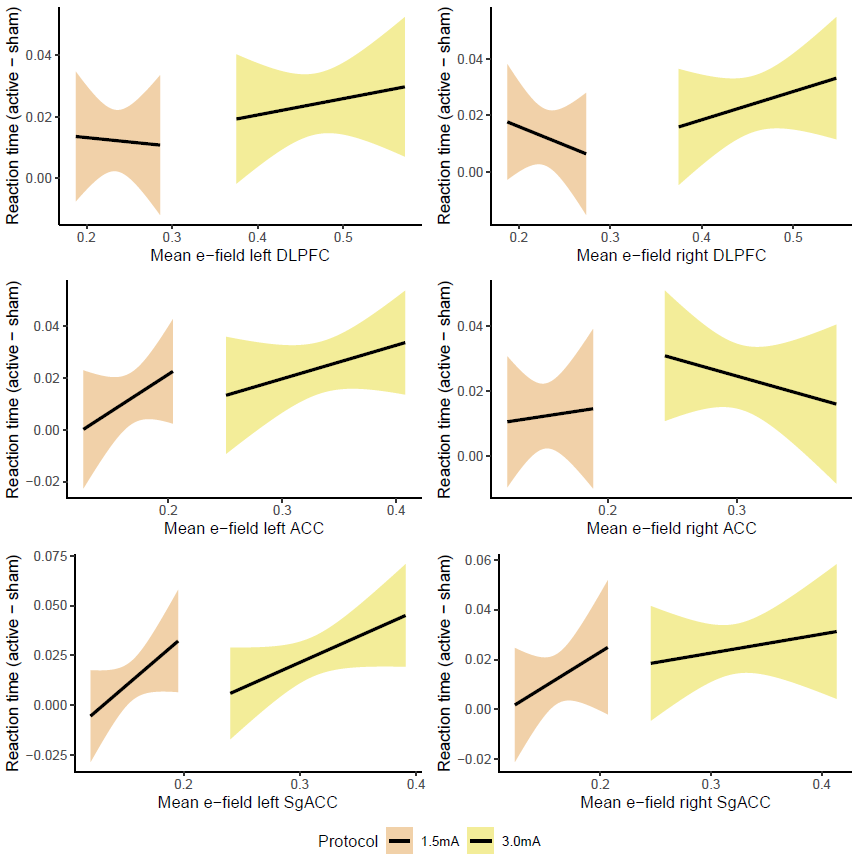

Abbreviation: ACC: Anterior Cingulate Cortex; DLPFC: Dorsolateral Prefrontal Cortex; SgACC: Subgenual Anterior Cingulate Cortex.

Visualization of the association between E-field in brain regions of interest and accuracy (active minus sham) in the emotional 3-task.

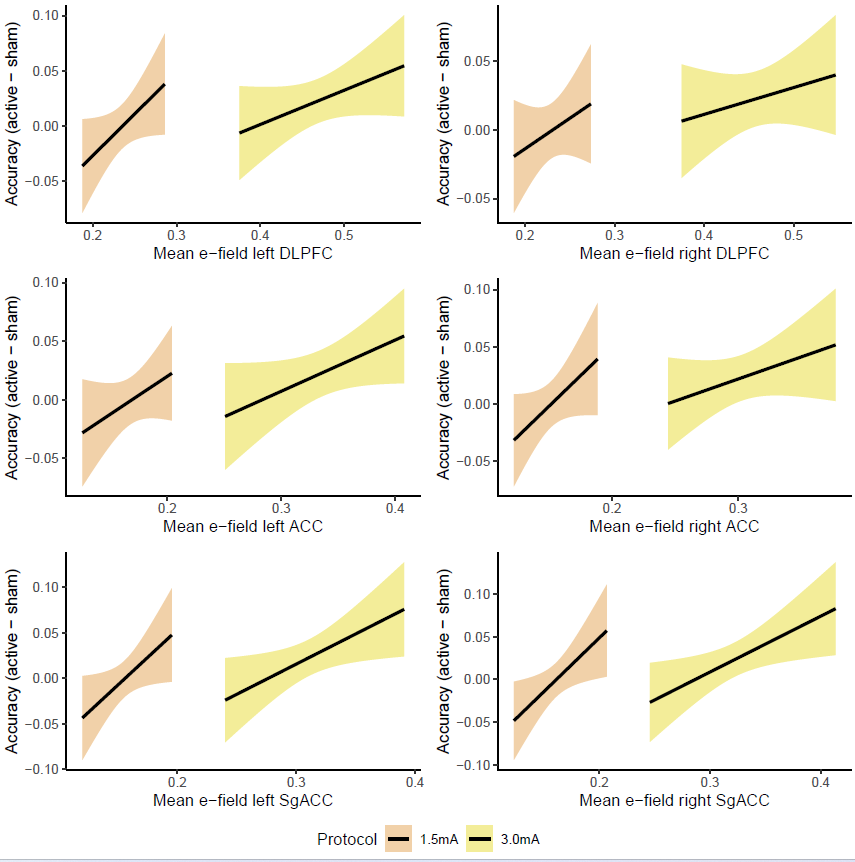

Abbreviation: ACC: Anterior Cingulate Cortex; DLPFC: Dorsolateral Prefrontal Cortex; SgACC: Subgenual Anterior Cingulate Cortex.

**Appendix 10**

**Table 1**. Frequence of adverse effects in each tDCS group

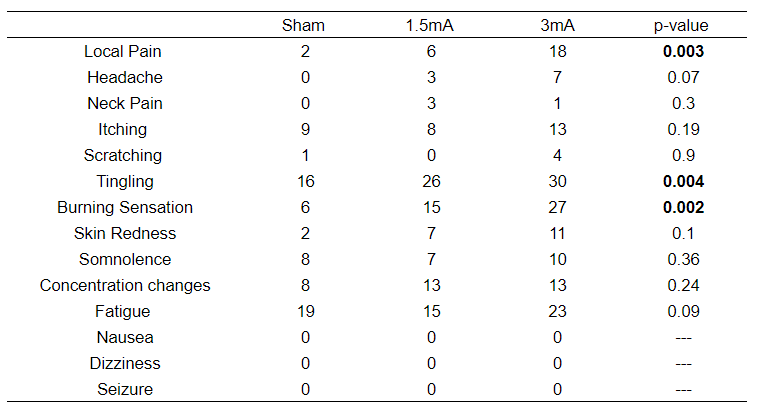
